# Offspring size resolves a population growth paradox in rays and skates

**DOI:** 10.1101/2024.01.02.573919

**Authors:** Ellen Barrowclift, Jennifer S. Bigman, Eric D. Digel, Per Berggren, Nicholas K. Dulvy

## Abstract

The maximum intrinsic population growth rate, *r*_max_, is a key determinant of the limits for sustainable fishing and is increasingly used in risk assessments. Metabolic theory suggests that *r*_max_ scales with adult body size, temperature (and hence depth) such that smaller-bodied species and those in warmer, shallower waters have greater *r*_max_ and, therefore, will be less sensitive to overexploitation. However, warm shallow-water tropical rays have lower *r*_max_ than cold deep-water temperate skates *contra* to the metabolic expectation. To resolve this paradox, we build from recent advances that suggest that offspring size may be key to understanding *r*_max_. Specifically, we examine how *r*_max_ is related to adult size, offspring size, temperature, and depth across 85 ray and skate species. Our results show that offspring size mediates relationships between *r*_max_, adult body size, temperature, and depth. Indeed, tropical rays had, on average, larger offspring and lower *r*_max_ compared to the temperate skates, despite living in warmer, shallower waters. Thus, despite the expectation from theory that tropical species should have faster life histories compared to temperate species, our result explains why tropical rays are actually less resilient. It remains unclear as to why tropical rays have such large offspring but we speculate that this is due to greater predation risk in shallow tropical waters driving the additional maternal investment in offspring size via the evolution of viviparity and matrotrophy. Our work highlights the complex relationships among life histories and the environment and may help explain global biogeographic patterns of intrinsic sensitivity to overexploitation.

## 1. Introduction

A key challenge in population biology is understanding global patterns of life histories, which is of both fundamental interest as well as useful for identifying species sensitivity to overfishing and other anthropogenic threats. Biogeographic patterns in life histories appear to be mediated by temperature (Beukhof *et al*., 2019), for example, Bergmann’s rule states that terrestrial endotherms in cooler environments will be larger-bodied than their warmer relatives (Bergmann, 1847). Similarly, the Temperature-Size Rule (TSR) describes the observed pattern that ectothermic populations and species generally grow faster to a smaller size at maturity (and presumably, smaller adult size) in warmer temperatures (Atkinson, 1994; Atkinson and Sibly, 1997; Atkinson, Morley and Hughes, 2006). Finally, metabolic theory suggests that species in warmer waters (e.g. in the tropics and shallow waters) with higher metabolic rates will tend to have ‘faster’ life histories than those in cooler waters (e.g. high latitude and deep waters) (Juan-Jordá *et al*., 2013; Beukhof *et al*., 2019; Wong, Bigman and Dulvy, 2021; Gravel *et al*., 2024). Typically, species with faster life histories grow faster to a smaller maximum body size, mature earlier, and have shorter lifespans, resulting in higher maximum intrinsic rate of population increase, *r*_max_ (Hutchings *et al*., 2012; Thorson, 2020; Gravel *et al*., 2024).

*r*_max_ is the average annual number of female spawners produced per female spawner at low population density (i.e. in the absence of density-dependence). Hence, *r*_max_ is an essential component of fisheries management to determine fishing limits and species’ recovery potentials (Cortés, 2016; Pardo *et al*., 2016; Horswill *et al*., 2025). There is considerable interest in calculating *r*_max_ for chondrichthyans but general patterns have proven hard to find (Mejía *et al*., 2025) unless trait data are carefully matched from similar habitats and consistent models are used (Pardo and Dulvy, 2022; Gravel *et al*., 2024; Moro *et al*., 2025). According to metabolic scaling expectations, *r*_max_ will scale with (adult) body mass (with an exponent of -0.25) and, independently, increase with temperature (Brown *et al*., 2004; Savage *et al*., 2004). This has been shown empirically both in experimentally-manipulated populations as well as across species in the wild (Luhring and Delong, 2017; Bernhardt, Sunday and O’Connor, 2018). For example, a positive relationship between *r*_max_ and temperature exists across Atlantic cod (*Gadus morhua*, Gadidae), which has greater *r*_max_ in warmer, more southerly populations (Myers, Mertz and Fowlow, 1997; Savage *et al*., 2004). Further, cooler, temperate tunas (Scombridae) with slower life histories experienced greater population declines than tropical lower-latitude tunas with faster life histories, after controlling for fishing mortality (Juan-Jordá *et al*., 2011, 2015). More generally, the ratio of production to biomass (P:B) changes systematically with latitude across the world’s fish communities, with high production and low standing biomass in the tropics compared to that found at cooler temperate and polar latitudes (Jennings *et al*., 2008). Recent work suggests that in addition to temperature and adult size, offspring size may also be an important determinant of *r*_max_ (Neuheimer *et al*., 2015; Denéchère, van Denderen and Andersen, 2022). This suggests that for two species with the same adult mass, one with larger offspring (and fewer of them) may have a lower *r*_max_ and therefore, offspring size could affect the scaling of *r*_max_ (Burger, Hou and Brown, 2019; Denéchère, van Denderen and Andersen, 2022).

Rays and skates of the superorder Batoidea are widely distributed across the world’s oceans but with a distinct biogeographic separation (Figure 1). Skates have an anti-tropical distribution and are typically found in the cooler waters of polar and temperate seas, as well as deeper, cool waters in the tropics, although there are also a few shallow-water tropical species (e.g. Teevan’s Skate *Dipturus teevani* and Rosette Skate *Leucoraja garmani*, Rajidae; Figure 1a). Whereas rays are generally found in the shallow tropics, subtropics and with seasonal occurrence in temperate waters during summer (Figure 1b), but with some deep-water exceptions (e.g. Deepwater Stingray *Plesiobatis daviesi*, Plesiobatidae, and Sixgill Stingray *Hexatrygon bickelli*, Hexatrygonidae) (McEachran and Miyake, 1990; Ebert and Compagno, 2007; Frisk, 2010). Metabolic theory suggests that warm-water rays should have higher *r*_max_, yet paradoxically, they have lower *r*_max_ than cold-water skates (Barrowclift *et al*., 2023). Most rays are live-bearers with very low fecundity and generally, have larger offspring (typically 1% of adult mass) compared to skates, which lay numerous eggs (mermaid’s purses) and overall, have smaller offspring (typically 0.1% of adult mass; Figure 2) (Goodwin, Dulvy and Reynolds, 2002; Mull *et al*., 2024). Here, we investigate whether the larger offspring size of tropical rays explains the paradox of their lower *r*_max_ compared to skates, while accounting for temperature and depth for 85 batoid species.

**Figure 1.**
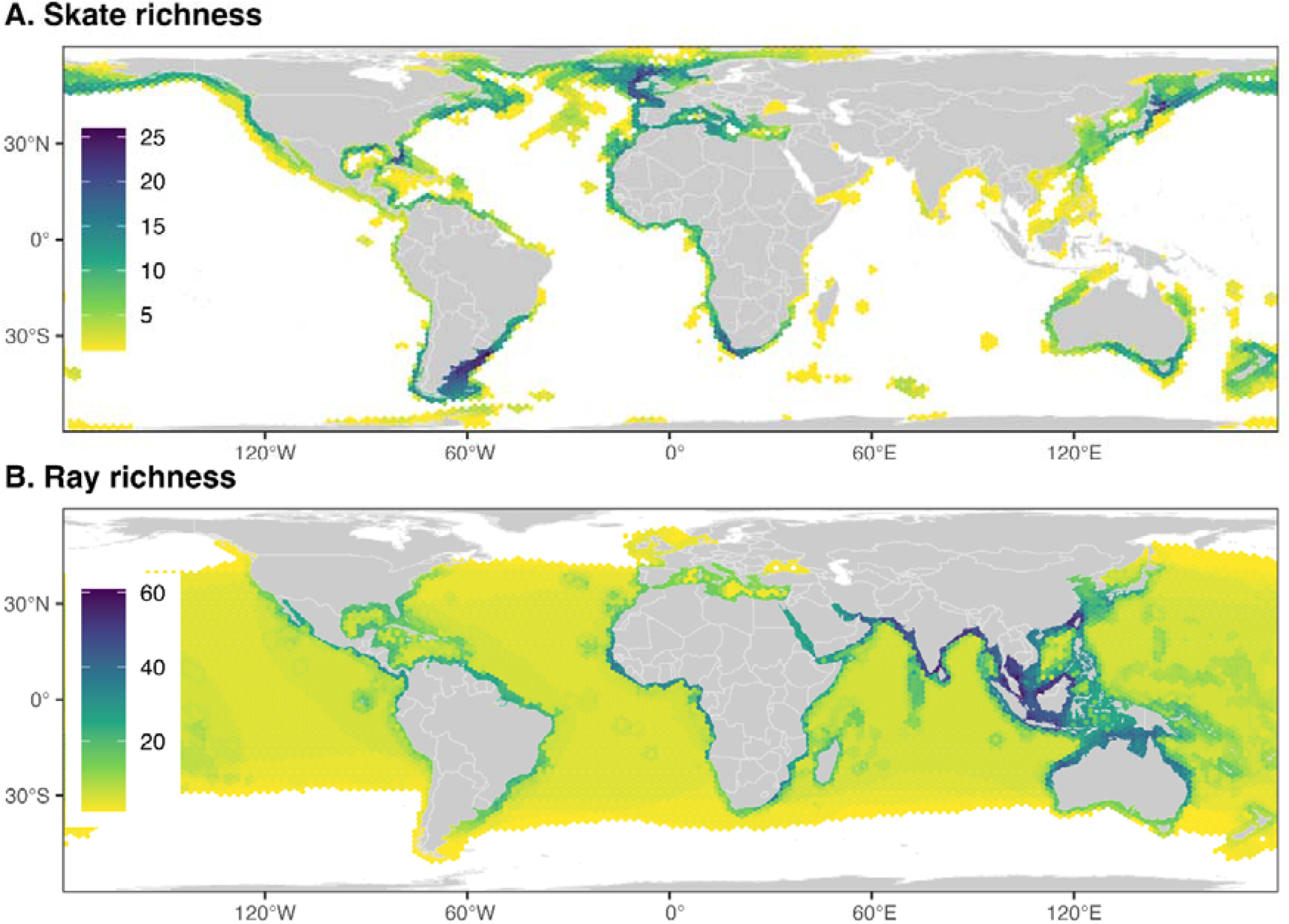
Biogeographic differences in the species richness of (A) skates (*n*=284 species of Rajiformes), which have a more anti-tropical distribution, compared to (B) rays, which have greater richness in the tropics (*n*=339 species of Torpediniformes, Rhinopristiformes and Myliobatiformes), mapped to a hexagonal grid with cell size of ∼ 23,322 km^2^.

**Figure 2:**
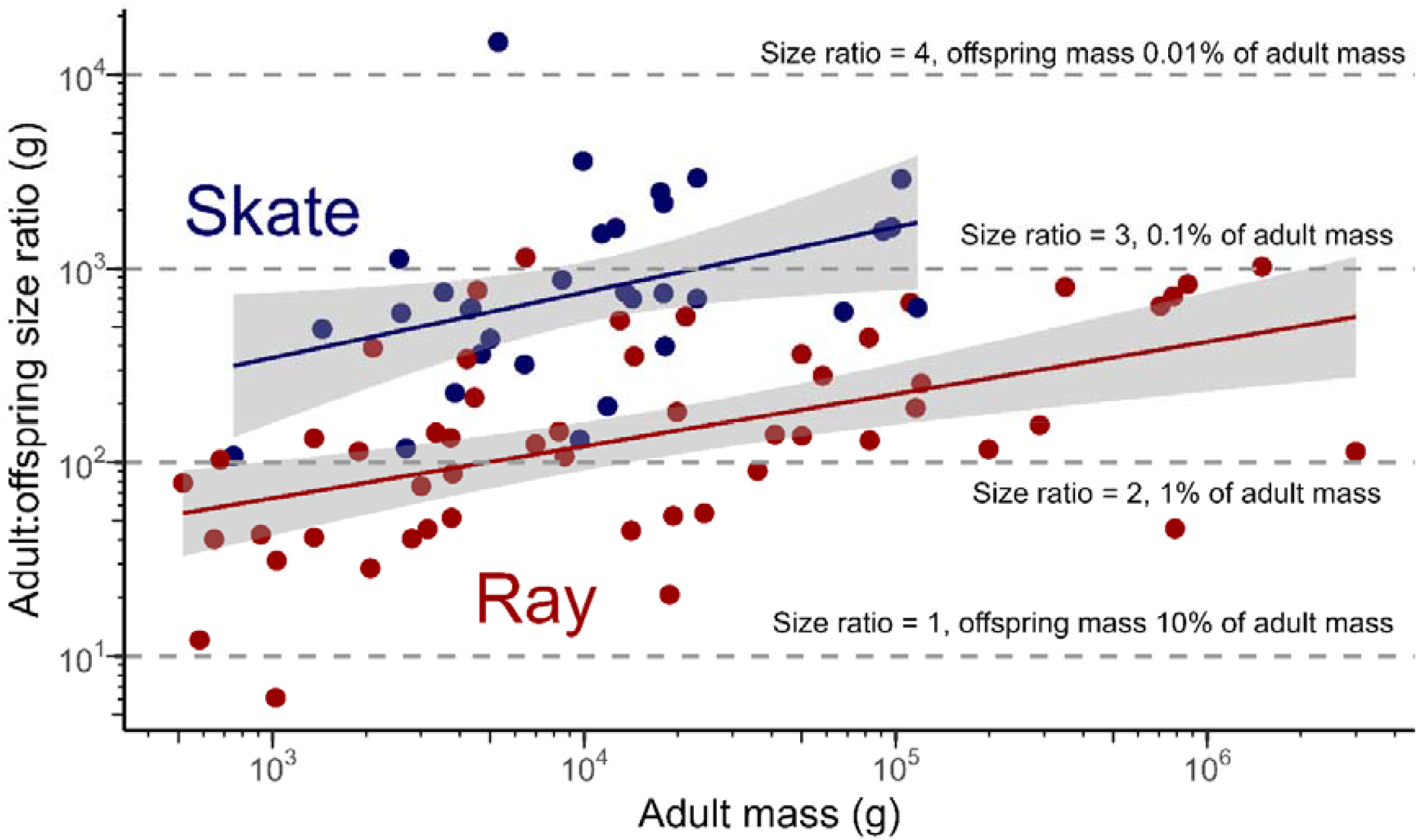
Relationship between adult to offspring size ratio (g) and adult mass (g) in log10 space for 85 rays (*n*=53, indicated by red points) and skates (*n*=32, indicated by blue points). The grey bands around the fitted models show the confidence intervals and grey dashed lines show adult to offspring size ratios of 1 to 4 where offspring size is 10 to 0.01% of adult body mass, respectively.

## 2. Material and Methods

We first describe the mapping of batoid species richness. Second, we present the calculation of *r*_max_, including the sources of the life history data used in the calculations. Third, we describe the calculation of environmental temperature-at-depth. Fourth, we summarise our analytical approach, including the statistical models used to assess different hypotheses of how *r*_max_ may vary with adult and offspring mass, temperature, and depth.

### 2.1 Mapping of ray and skate richness

We downloaded species distribution maps from the IUCN Red List as shapefiles (https://www.iucnredlist.org/resources/spatial-data-download). These were mapped to a hexagonal grid with cell sizes averaging 23,322 km^2^ and species richness was calculated per cell for each of skate (Order Rajiformes) and ray species (Orders: Torpediniformes; Rhinopristiformes, and Myliobatiformes) (Dulvy *et al*., 2014, 2021).

### 2.2 The calculation of *r*_*max*_ and source of life history data

The maximum intrinsic rate of population increase, *r*_max_, was calculated using a modified Euler-Lotka model, with a mortality estimator that accounts for survival to maturity with the following equation:

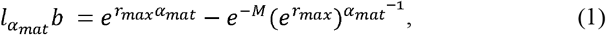

where *l*_*αmat*_ is the proportion of individuals surviving to maturity, which is calculated from annual fecundity *b* and the species-specific instantaneous natural mortality rate *M* (Cortés, 2016; Pardo *et al*., 2016). Natural mortality (*M*) was estimated as *M* = 1/*ω* (Dulvy et al., 2004) where w is an estimate of average lifespan in years and was assumed to be the midpoint between age at maturity (α_*mat*_) and maximum age (α_*max*_) (Pardo *et al*., 2016). The life history traits were sourced for 85 ray (Torpedo rays, Torpediniformes; Rhino rays, Rhinopristiformes; and stingrays, Myliobatiformes; *n*=53) and skate species (Rajiformes; *n*=32) (Figure 3). Three traits were geographically matched where possible to minimise spurious errors due to intraspecific life history variation (Gravel *et al*., 2024; Moro *et al*., 2025): 1) female age at 50% maturity (α_*mat*_; years), 2) maximum age for females, where known (α_*mat*_; years), and 3) annual reproductive output (number of female offspring assuming 1:1 sex ratio; *b*) reported in a published global life history database (Barrowclift and Dulvy, 2023) compiled in Barrowclift et al. (2023).

**Figure 3.**
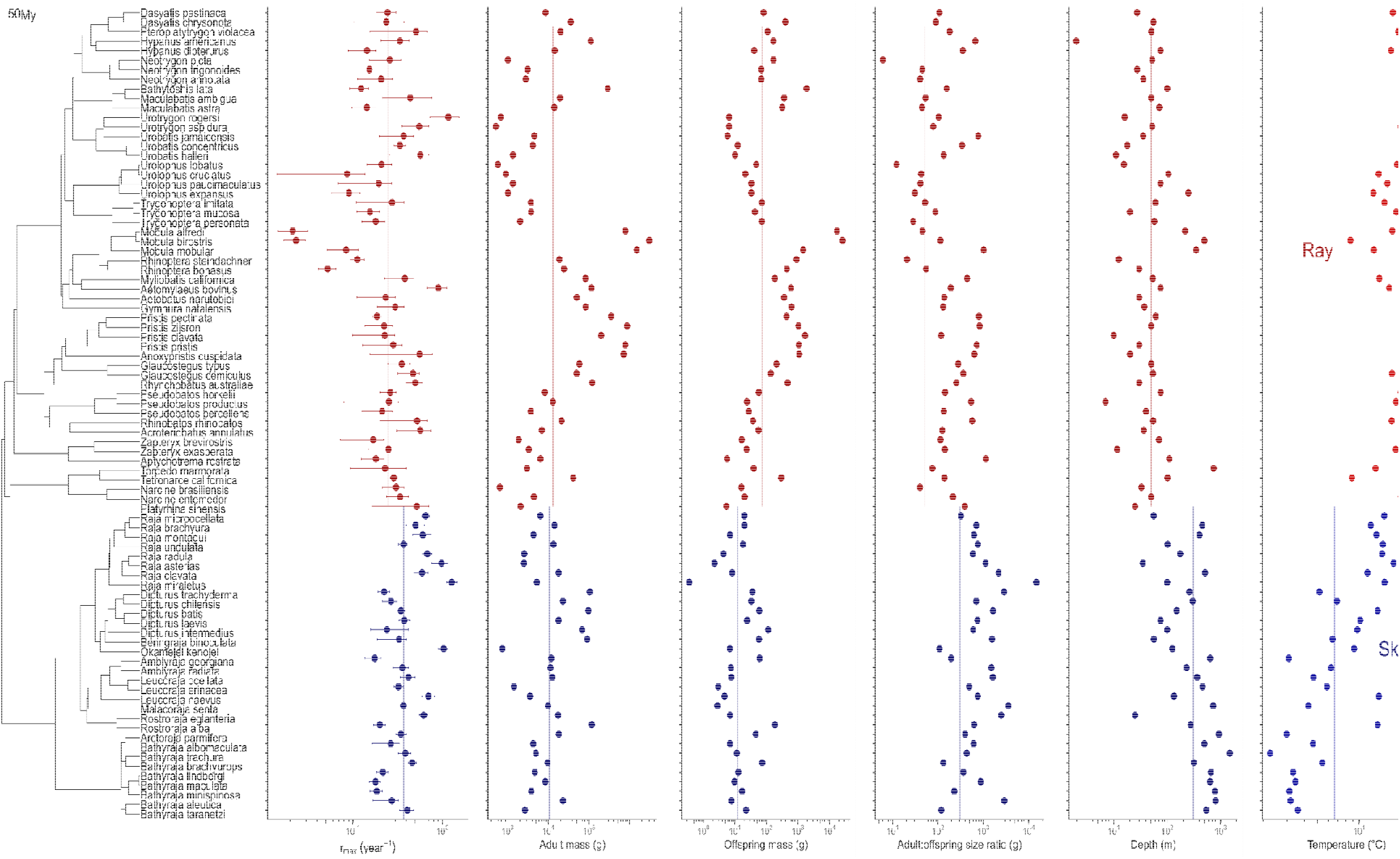
Phylogeny, maximum intrinsic rate of population increase (*r*_max_), adult and offspring mass (g), offspring to adult size ratio, depth (m), temperature (^◦^C) in log10 space for 85 rays (*n*=53, red points) and skates (*n*=32, blue points). Solid lines show median values. Uncertainty in *r*_max_ estim shown with 2.5% and 97.5% quantiles. Phylogenetic tree from Stein et al. (2018) (available on Vertlife.org) with binomial nomenclature updated to ref current taxonomic nomenclat

Adult body mass (maximum weight in grams) for the 85 batoid species was also sourced from the published life history database compiled in Barrowclift et al. (2023). Offspring mass (in grams) was estimated from offspring size measured as total length for skates or disc width for rays in centimetres (cm) as reported in Rays of the World (Last *et al*., 2016), Sharkipedia (Mull *et al*., 2022), and IUCN Red List assessments (Dulvy *et al*., 2021). Offspring size was not available for 14 species and for these cases we estimated offspring size from related species with similar maximum length and body shape (Barrowclift and Dulvy, 2023). The median offspring length was calculated in cases when minimum and maximum offspring lengths were reported. The corresponding offspring mass was then calculated using length-weight regression coefficients extracted from FishBase using the package *rfishbase* (Boettiger, Lang and Wainwright, 2012; Froese and Pauly, 2022; Barrowclift and Dulvy, 2023). Length-weight regression coefficients were selected for females where possible. When length-weight regressions were not available, we used parameters for a closely related species with similar body shape and maximum size (Barrowclift and Dulvy, 2023).

### 2.3 Calculation of environmental temperature-at-depth

Median depth and environmental temperature for the 85 ray and skate species were also taken from Barrowclift et al. (2023). Specifically, median depth estimates for each species were taken from depth ranges of IUCN Red List assessments as compiled in Dulvy et al. (2021). Temperature-at-depth data were determined by intersecting each species distribution map with the International Pacific Research Center’s interpolated dataset of gridded mean annual ocean temperatures. This dataset is based on measurements from the Argo Project (data available at http://apdrc.soest.hawaii.edu/projects/Argo/data/statistics/On_standard_levels/Ensemble_mean/1x1/m00/index.html). Species range maps were sourced from https://www.iucnredlist.org/resources/spatial-data-download. Temperature grid points were extracted across the species’ distribution from the depth level that was closest to the species’ median depth and the median temperature calculated (Pardo and Dulvy, 2022; Barrowclift *et al*., 2023).

### 2.4 Statistical inference

Metabolic scaling expectations for how *r*_max_ relates to adult body mass and temperature (Savage *et al*., 2004) can be estimated with the following linear model:

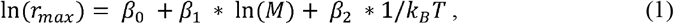

where *r*_*max*_ is the maximum intrinsic rate of population increase (year^-1^), *β*_0_ is the intercept, *β*_1_ is the body mass-scaling coefficient, *M* is adult mass in grams, *β*_2_ is the activation energy *E, T* is the temperature (in Kelvin), and *k*_*B*_ is the Boltzmann constant (8.617 × 10^-5^ eV).

We modified this initial equation (1) above to test whether absolute offspring size (*M*_*offspring*_) and the adult to offspring mass ratio (*M*/*M*_*offspring*_) explained variation in *r*_max_ across species in addition to temperature and depth (Denéchère, van Denderen and Andersen, 2022). *M*/*M*_*offspring*_ was suggested by Denéchère et al., (2022) to be a better predictor of *r*_max_ than either adult or offspring body mass and thus it is examined here. Specifically, we used an information-theoretic approach to compare a total of 32 models that represented different hypotheses as to how *r*_max_ may vary across species (see Table S1 for hypotheses, models, and AIC values and weights).

For model fitting, *r*_max_, adult and offspring masses were natural log-transformed, and temperature and depth data were standardised (scaled and centred). We found that offspring body mass and adult body mass (Figure 4a) as well as temperature and depth were collinear (Pearson’s r > 0.75), but there was no relationship between either adult or offspring body mass and temperature (Figure 5). Thus, we only included either offspring body mass or adult body mass and either temperature or depth in a given model, and did not consider interactions between either mass and either temperature or depth. All models were run in a phylogenetic comparative framework to account for the shared evolutionary history between species. To do so, we fit phylogenetic generalised linear models using the *pgls* function in the *caper* package (Orme *et al*., 2018). Phylogeny was used as a random effect in all models using a phylogenetic tree randomly drawn from the distribution of chondrichthyan trees (Stein *et al*., 2018) (available at Vertlife.org; Figure 3). The binomial nomenclature of the trees was updated to reflect current taxonomic nomenclature. The phylogenetic position of two species was not known (*Aetobatus narutobiei*, Aetobatidae, and *Maculabatis ambigua*, Dasyatidae), and therefore, two closely related species (*A. flagellum* and *M. gerrardi*, respectively) were used instead. Models were refitted with an additional 10 random trees to test the sensitivity of results to slight variations in the phylogenies; temperature and depth were interchangeable in the top model, and therefore, results were reported for a single tree (Table S2 in the Supporting Information). The corrected Akaike Information Criterion (AICc) were used to compare models. If including a parameter improved the model’s AICc by less than two units (ΔAICc ≤ 2), it was considered relatively uninformative (Burnham and Anderson, 2002; Arnold, 2010). All analyses were run in R version 4.1.2 (R Core Team, 2021) in RStudio (RStudio Team, 2021).

**Figure 4.**
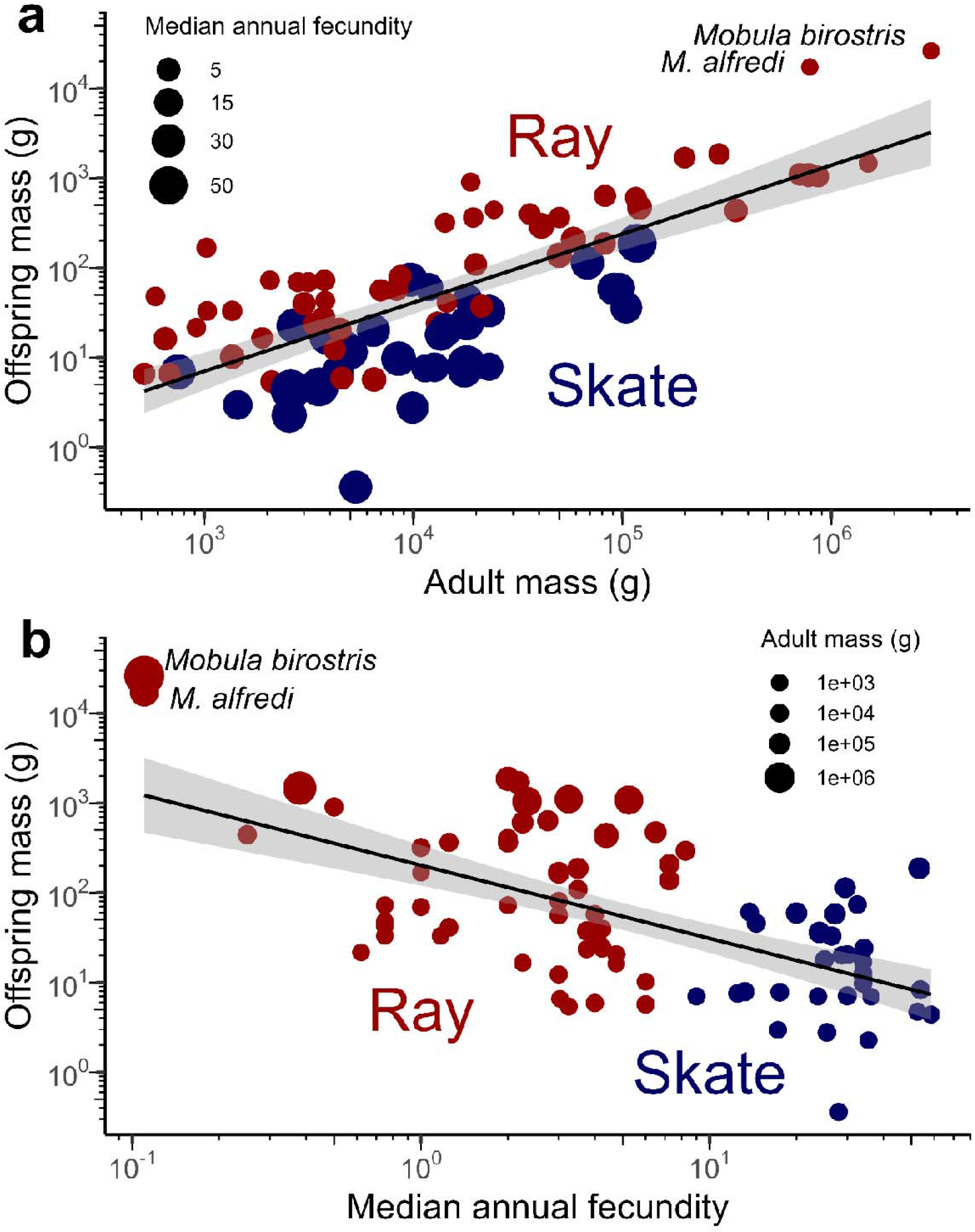
Relationships between offspring body mass and (a) adult body mass (with point size showing median annual fecundity) and (b) median annual fecundity (with point size showing adult mass) on a log10 scale for 53 rays (red points) and 32 skates (blue points).

**Figure 5:**
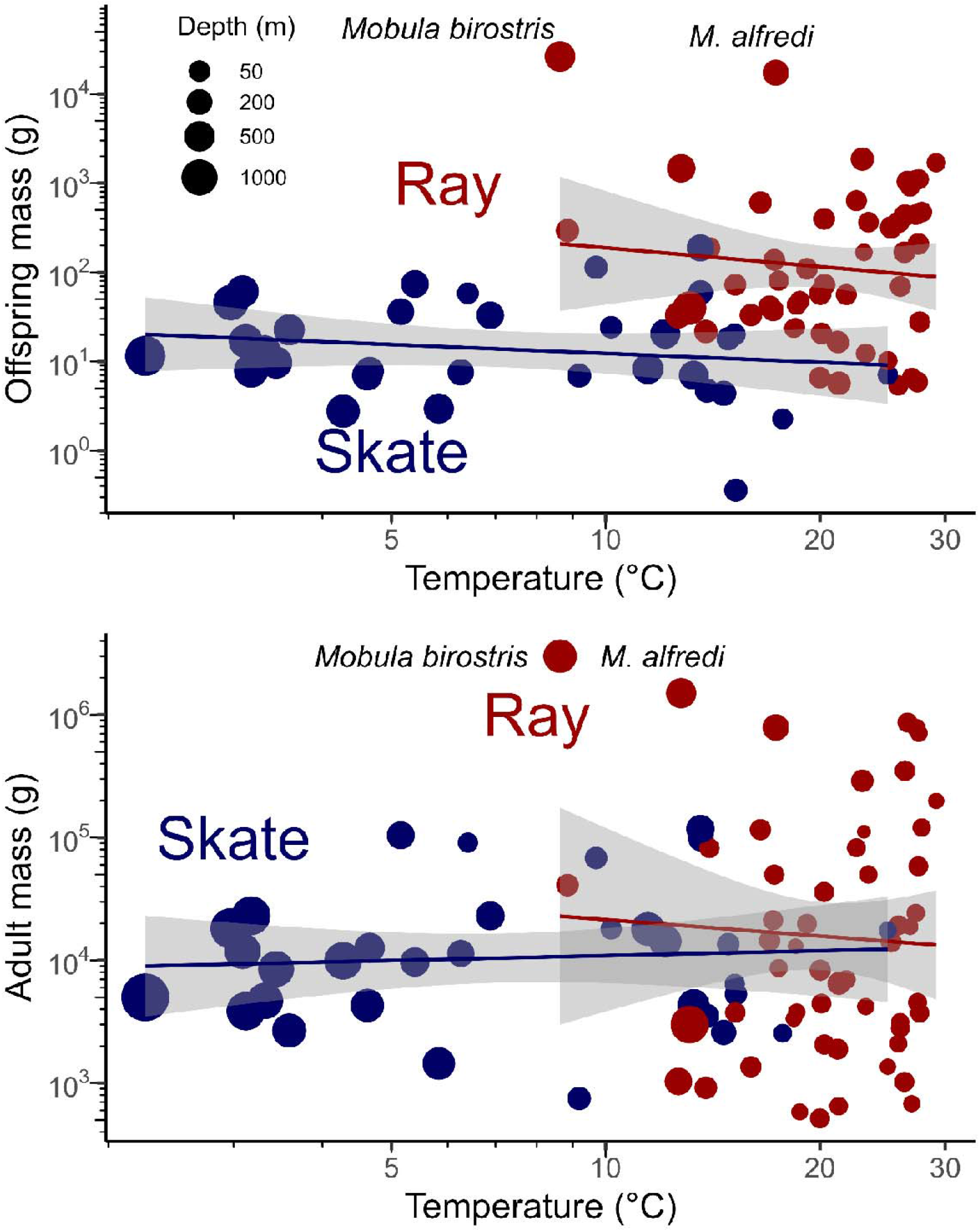
Relationship between temperature (^◦^C) and a) offspring body mass (g) and b) adult body mass (g) in log space for 85 rays (*n*=53, indicated by red points) and skates (*n*=32, indicated by blue points). Median depth of occurrence (m) is shown by the point size, with a linear model fitted to ray and skate data points. The grey bands around the fitted models show the confidence intervals.

## 3. Results

Tropical rays had, on average, larger offspring and lower *r*_max_ compared to the temperate skates, despite living in warmer and shallower waters. We found that the ratio of adult to offspring size was not important in explaining variation in *r*_max_ and thus, we focus on the 11 models without this ratio in the main text (see Table S1 in the Supporting Information for results of the models with the adult to offspring size ratio). Our findings were insensitive to the exclusion of the two species with the largest offspring masses: *Mobula alfredi* and *M. birostris* (Mobulidae). The top models were the same and therefore results were presented for the full 85 species dataset (Table S3 in the Supporting Information).

Offspring size combined with either temperature or depth were the most important variables explaining the global variation in *r*_max_ across batoids (Figures 6 and 7). Of the 11 models examined, the most supported model suggests that *r*_max_ varies with offspring mass and depth (ΔAICc = 0), describing the greatest amount of variation in *r*_max_ across species (adjusted *R*^*2*^ = 0.20; Table 1). The second-ranked model described *r*_max_ varying with offspring mass and temperature (ΔAICc = 0.3) and had approximately 86% of the support of the top-ranked model (when compare the Akaike weights) and accounted for slightly less variation (adjusted *R*^*2*^ = 0.19; Table 1). These top two models – one with depth and one with temperature – were essentially equivalent in terms of support suggesting that either variable could be combined with offspring mass to explain the variation in *r*_max_ across rays and skates (Figure 6 and 7). Whether including temperature or depth as statistically the top model depended on the phylogeny (Table S2). In all models, the effect sizes of offspring mass, depth, and temperature had 95% confidence intervals that did not overlap zero (Figure 6). A model describing *r*_max_ varying solely with offspring mass received moderate support but with only 22% of the support of the top-ranked model and accounting for less variation (adjusted *R*^*2*^ = 0.16). Therefore, including temperature or depth in the model improved the model support (based on AIC comparison; Table 1).

**Table 1:** The 11 models examined with associated hypotheses for how maximum intrinsic rate of population increase (*r*_max_) varies with offspring body mass (*M*_*offspring*_), adult body mass *M*, inverse temperature 1/*k*_*B*_*T*, and depth. Comparison of models using corrected Akaike Information Criteria (AICc), number of parameters (n), negative log-likelihood (-LL), adjusted R^2^, difference in AICc from the top model (ΔAICc), and Akaike weights. Models are ordered by ascending AICc, with models with AICc < 2 shown in bold.

| Hypothesis | Model: $\ln(r_{\max}) \sim$ | n | -LL | AICc | $R^2$ | Adj. $R^2$ | $\Delta\text{AICc}$ | Weights |
| --- | --- | --- | --- | --- | --- | --- | --- | --- |
| $r_{\max}$ varies with offspring body mass and depth | $\ln(M_{\text{offspring}}) + \text{depth}$ | **3** | **-62.8** | **132** | **0.22** | **0.2** | **0** | **0.451** |
| $r_{\max}$ varies with offspring body mass and temperature | $\ln(M_{\text{offspring}}) + 1/k_B T$ | **3** | **-63** | **132.3** | **0.21** | **0.19** | **0.3** | **0.388** |
| $r_{\max}$ varies with offspring body mass only | $\ln(M_{\text{offspring}})$ | 2 | -65.4 | 135 | 0.17 | 0.16 | 3 | 0.101 |
| $r_{\max}$ varies with adult body mass and temperature | $\ln(M) + 1/k_B T$ | 3 | -66.1 | 138.5 | 0.14 | 0.12 | 6.5 | 0.017 |
| $r_{\max}$ varies with adult body mass and depth | $\ln(M) + \text{depth}$ | 3 | -66.1 | 138.6 | 0.14 | 0.12 | 6.6 | 0.017 |
| $r_{\max}$ varies with adult body mass and temperature, and the effect of mass scaling coefficient varies with temperature | $\ln(M) * 1/k_B T$ | 4 | -65.7 | 139.8 | 0.16 | 0.13 | 7.8 | 0.009 |
| $r_{\max}$ varies with adult body mass only | $\ln(M)$ | 2 | -68 | 140.2 | 0.1 | 0.09 | 8.2 | 0.007 |
| $r_{\max}$ varies with adult body mass and depth, and the effect of mass scaling coefficient varies with depth | $\ln(M) * \text{depth}$ | 4 | -66.1 | 140.8 | 0.14 | 0.11 | 8.8 | 0.006 |
| $r_{\max}$ varies with (inverse) temperature only | $1/k_B T$ | 2 | -69.5 | 143.1 | 0.07 | 0.06 | 11.1 | 0.002 |
| $r_{\max}$ varies with depth only | depth | 2 | -70.3 | 144.8 | 0.05 | 0.04 | 12.8 | 0.001 |
| average $r_{\max}$ (i.e. intercept-only model) | 1 | 1 | -72.7 | 147.5 | 0 | 0 | 15.5 | 0 |

**Figure 6:**
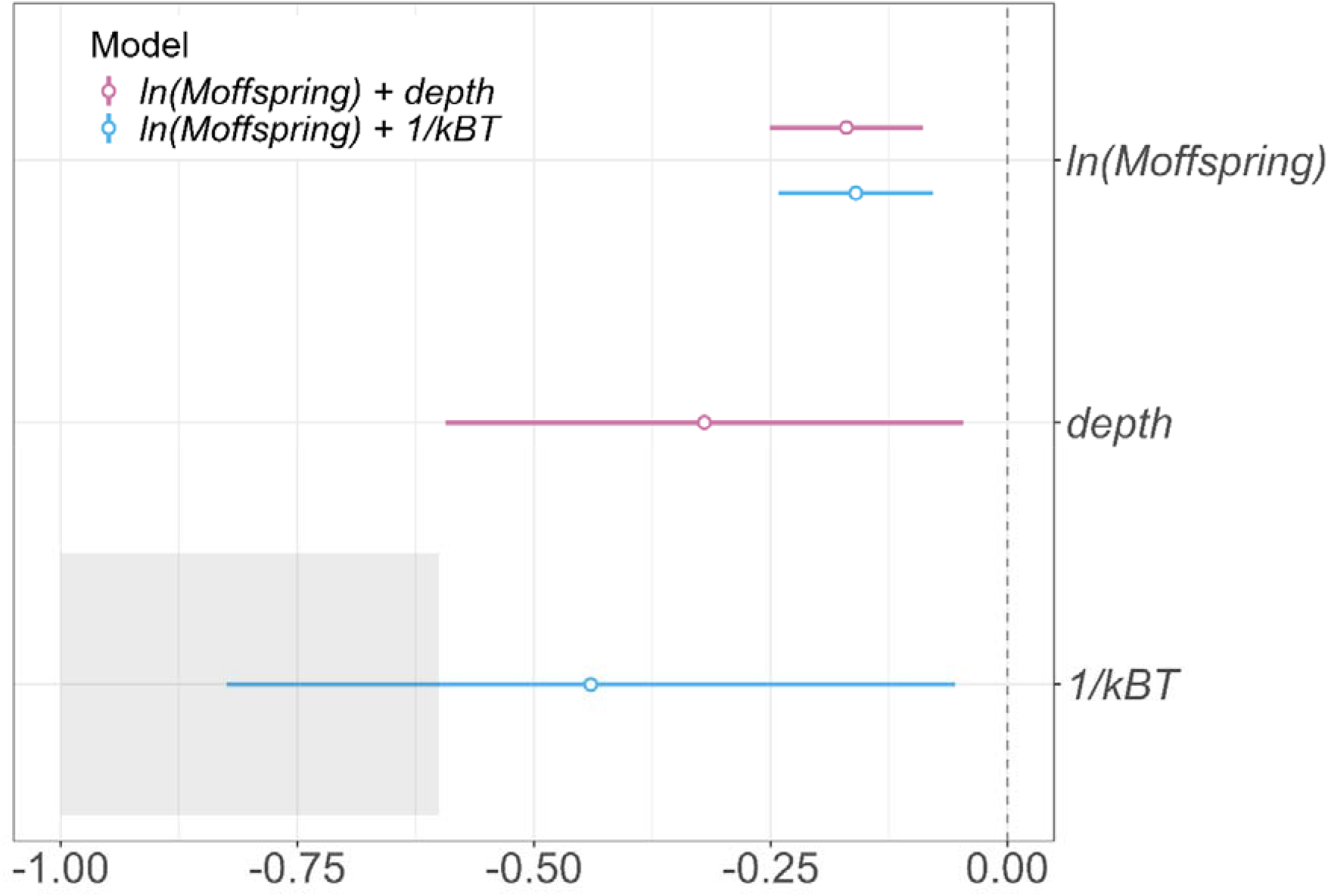
Coefficient plot showing the effect sizes for offspring body mass (*M*_*offspring*_), depth, and inverse temperature (1/*k*_*B*_*T*) on *r*_max_ in the top two models. Error bars show the 95% confidence intervals and effect sizes are considered significant when confidence intervals do not overlap zero. The grey shaded box shows the expected effect size for temperature (-1.0 to -0.6) based on metabolic theory.

**Figure 7:**
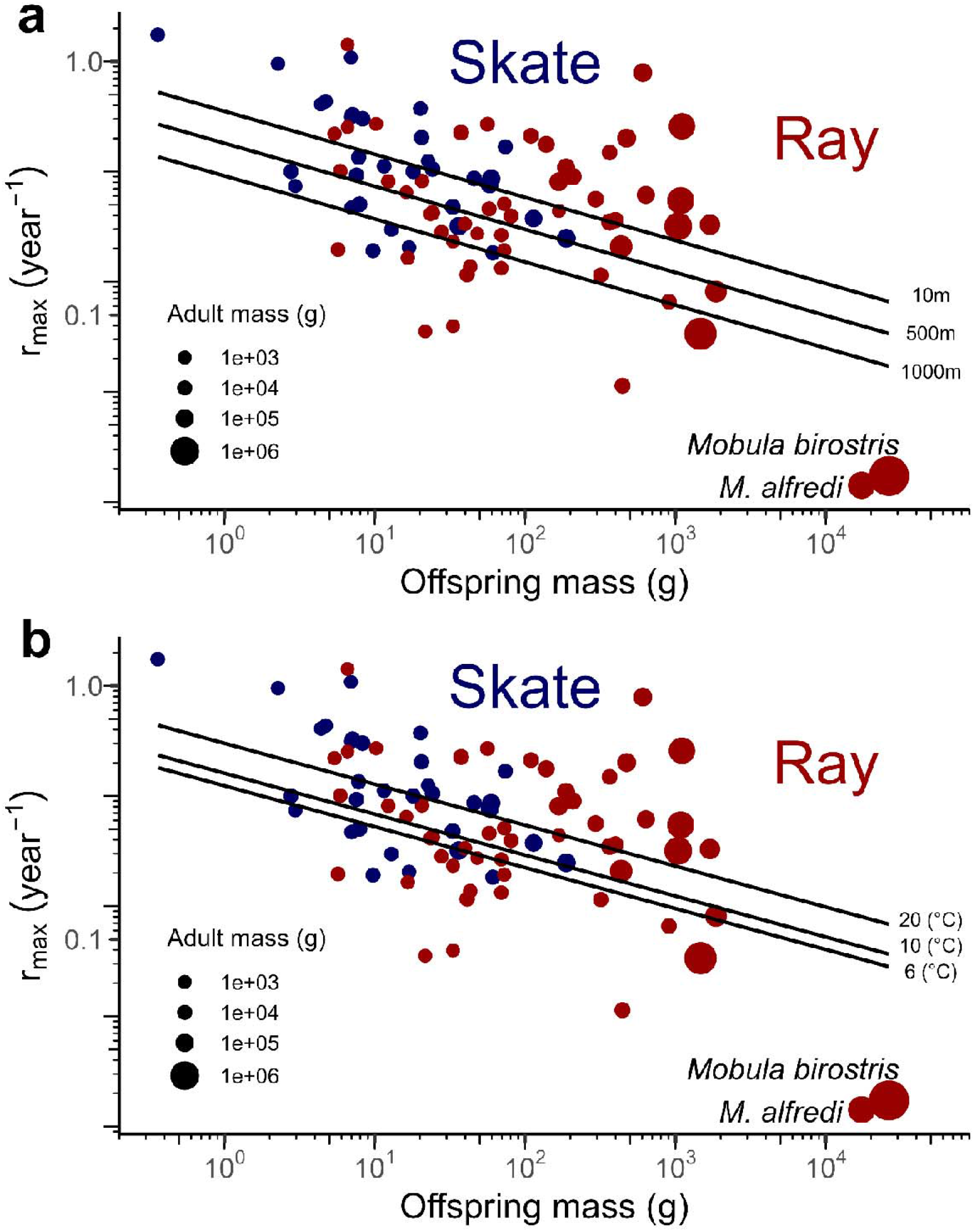
Relationship between maximum intrinsic rate of population increase (*r*_max_) and offspring body mass, showing the effects of either (a) depth or (b) environmental temperature for 85 rays (*n*=53, red points) and skates (*n*=32, blue points) in log10 space. Fitted lines show the predicted relationships for the two top models: (a) ln(*r*_max_) ∼ ln(*M*_*offspring*_) + depth for three depths (10, 500, 1000m) and (b) ln(*r*_max_) ∼ ln(*M*_*offspring*_) + for three temperatures (6, 10, 20°C). Adult body mass is shown by the point size.

The effect of offspring mass on maximum population growth rate, *r*_max_, of batoids is negative, indicating that *r*_max_ is lower in species with larger offspring sizes and higher in species with smaller offspring sizes (Figure 7; Table S4 in the Supporting Information). The effect of inverse temperature 1/*k*_*B*_*T* and depth were negative across models indicating *r*_max_ is higher in warmer-shallower water species as would be expected from metabolic theory (Figure 7; Table S4). The scaling of temperature with *r*_max_ overlapped with the expectation of approximately -0.6 from metabolic theory (Figure 6; Table S4). *r*_max_ was greater for species at shallower depths (Figure 7a) or at warmer temperatures (Figure 7b) based on the fitted top two models for the predicted relationships across three depths and three temperatures, respectively.

By comparison, the remaining models were not well supported (ΔAICc > 6; Table 1). Adult mass did not have as much support as offspring mass in explaining variation in *r*_max_ across rays and skates and models without any measure of size were far less supported than those with either offspring or adult size (Table 1). Indeed, the effect of adult mass on *r*_max_ was approximately half (-0.12) than expected from metabolic theory (-0.25; Table S4 in the Supporting Information).

For all models, there was a strong phylogenetic signal from the residuals of *r*_max_ (Pagel’s *λ* ≥ 0.79; Table S4 in the Supporting Information).

## 4. Discussion

We find that offspring size disrupted the well-studied relationships between *r*_max_, (adult) body size, and temperature across 85 ray and skate species. Specifically, our results show that temperature (or depth) and adult body size alone – as previously thought – do not explain the variation in *r*_max_ across rays and skates. Instead, offspring size appears to be the key predictor of *r*_max_ in these taxa such that species with larger absolute offspring size have lower population growth rates, *r*_max_. This finding helps explain the paradox of why tropical rays, which are found in warmer, shallower waters and should have higher *r*_max_ based on expectations from metabolic and life history theory, instead have lower population growth rates, compared to cold, deep-water skates (which have smaller offspring size).

Life history and metabolic theory suggest that in warmer temperatures, organisms generally grow faster, mature earlier, and attain smaller maximum body sizes resulting in faster generation times and higher production to biomass ratios (Beukhof et al., 2019; Jennings et al., 2008; Munch & Salinas, 2009). This would also lead to latitudinal and depth-related patterns of temperature and size, whereby organisms in shallower, warmer, and/or lower latitude waters have higher metabolic rates and therefore ‘faster’ life histories compared to organisms in deeper, cooler, and/or higher latitude waters (Juan-Jordá *et al*., 2013; Wong, Bigman and Dulvy, 2021; Pardo and Dulvy, 2022; Gravel *et al*., 2024). However, recent work found that compared to the cold water, deeper-dwelling skates, warm, shallow-water tropical rays have lower *r*_max_ and therefore greater intrinsic sensitivity to anthropogenic threats such as overfishing (Barrowclift *et al*., 2023). Here, we delve into this paradox and uncover that offspring size modulates how *r*_max_ scales with temperature (or depth) and adult body size. Specifically, offspring size, and not adult size, was the most important size metric for explaining variation in *r*_max_ across batoid species and along with temperature or depth, explained the most variation. Generally, offspring size tends to have a negative relationship with temperature in vertebrates and invertebrates due to differences in maternal investment, with females producing larger, better-provisioned offspring in colder environments (Pettersen *et al*., 2020; Marshall, 2021). This large-offspring-in-the-cold pattern is supported by both a cross cohort experiment of bryozoans (Marshall, 2021) and a metanalysis spanning 72 species from five ectotherm phyla (Pettersen *et al*., 2020). However, tropical rays generally have larger offspring sizes than cooler-water-temperate skates. It may be that offspring size is largely independent of temperature for batoids, with both employing very different reproductive strategies, raising the question as to why large offspring sizes have evolved in shallow-water, tropical rays. Collectively, our work suggests that at least for some taxa, the pattern of reproductive investment into offspring size can interfere with the effect temperature and size typically has on other life history traits.

Our finding that offspring size explains variation in *r*_max_ across batoids follows from recent work suggesting that the scaling of *r*_max_ with adult body mass matches metabolic expectations only when offspring size is considered, and specifically, when offspring mass is proportional to adult mass (constant adult-to-offspring size ratio) (Denéchère, van Denderen and Andersen, 2022). For elasmobranchs, Denéchère *et al*. (2022) found that the scaling of *r*_max_ with adult mass matched metabolic expectations and that elasmobranchs had a constant adult-to-offspring ratio (the ratio of adult size to offspring size does not increase with adult mass). We found that the ratio of adult size to offspring size was not important in the scaling of *r*_max_ and that in batoids, this ratio increases with adult size (i.e. it is not constant as found in Denéchère *et al*. (2022) when examined across a broader diversity of elasmobranchs). Further, we found that the effect size of absolute offspring mass on *r*_max_ was larger than adult mass, suggesting that an increase in offspring size limits *r*_max_ more strongly than adult size, and that using the ratio of adult size to offspring size had low predictive power. Offspring size, which reflects reproductive investment and life history trade-offs, greatly varied across the 85 species we examined, which may be why we were able to detect the importance of this trait. Specifically, the differences in *r*_max_ between warm, shallow-water rays and cold water, deeper-dwelling skates are likely due to their different reproductive strategies – as live-bearing rays have fewer, larger offspring compared to egg-laying skates with large numbers of smaller offspring – as hypothesised in Barrowclift *et al*. (2023).

Live-bearing has been hypothesised to have evolved from egg-laying in order to increase the survival of offspring through a controlled maternal environment and greater protection from predators (Wourms, 1994; Goodwin, Dulvy and Reynolds, 2002). In live-bearing species, offspring size is constrained by the size of the maternal body cavity but results in offspring with greater survival (Wourms and Lombardi, 1992; Musick and Ellis, 2005). Whereas in egg-laying species, size is limited by nutrients stored in the yolk sac (Carrier, Pratt and Castro, 2004). The egg-laying reproductive strategy of skates is thought to be advantageous because it requires less energy and shorter reproductive cycles but there will be survival consequences for the offspring due to smaller size, and, thus, greater risk of predation (Goodwin, Dulvy and Reynolds, 2002). Consequently, there is a trade-off between the number and size of offspring resulting in a negative relationship between offspring size and annual fecundity (Smith and Fretwell., 1974; Duarte and Alcaraz, 1989; Cortés, 2000). This was evident in our batoid dataset along with the expectation that larger species tend to have larger and more offspring (Cortés, 2000). Offspring size will affect juvenile survival to maturity, which is important to consider, given the maternal trade-off between lifetime reproductive output, which likely varies between egg-laying and live-bearing reproductive modes (Pardo *et al*., 2016). In the calculation of *r*_max_ for elasmobranchs, there is the pragmatic assumption that juvenile survival to maturity is the same as the survival rate of adult ages (a consistent mortality estimator). However, our key finding suggests the average mortality depends on offspring size, presumably with larger absolute offspring sizes (typical of tropical rays) having lower predation risk than smaller offspring (typical of colder-water skates). This then leads to the question as to why it might be advantageous for tropical, warm-water rays to have larger offspring than temperate, cold-habitat skates?

The ancestral reproductive mode of sharks and rays is egg-laying with the subsequent evolution of live-bearing and a particularly high degree of maternal investment found in the shallow-tropical elasmobranch species (Dulvy and Reynolds, 1997; Blackburn and Hughes, 2024; Mull *et al*., 2024). The diversification and radiation of elasmobranchs throughout shallow tropical shelf seas and the pelagic zone appears to be associated with the evolution of live-bearing and multiple mechanisms for providing additional maternal investment in offspring (Mull *et al*., 2024). The question remains as to why live-bearing with additional maternal investment has evolved. Predation tends to be size-based in the marine realm (Verity and Smetacek, 1996; Barnes *et al*., 2010). We speculate that shallow-water, tropical rays have evolved larger offspring in response to selection pressure from greater predation risk in the tropics. Increased offspring size reduces the threat of predation in systems structured by size-based interactions, i.e. has been selected to reduce juvenile mortality (Cortés, 2000; Olsson, Gislason and Andersen, 2016; Sibly *et al*., 2018). We speculate that this predation risk drove the evolution of live-bearing, and in particular the convergent evolution of multiple forms of matrotrophy (maternal supply of nutrients during gestation). Generally, offspring size is larger in chondrichthyans compared to teleost fishes in which live-bearing appears to have evolved in particularly small-bodied taxa, suggesting the drivers of viviparity are fundamentally different in chondrichthyans (Goodwin, Dulvy and Reynolds, 2002). Predation risk has generally been hypothesised to increase towards the tropics, with recent empirical work finding greater predation rates on epifaunal communities in shallow, tropical waters compared to high latitude waters (Ashton *et al*., 2022). This is relevant to potential greater predation on eggs and juveniles in tropical waters. Fisheries-driven decline in sharks (which predate on batoids) has led to increases in ray abundance, consistent with our hypothesis that predation is a major evolutionary force in tropical systems (Sherman *et al*., 2020; Simpfendorfer *et al*., 2023).

Given Bergmann’s rule and the Temperature-Size-Rule (TSR), adult body size of temperate skates in cooler waters would be expected to be larger than tropical rays in warmer waters. However, our results suggest there is wide variation in adult body size across batoids and, generally, the tropical rays are larger than cooler-water skates. We speculate above that the larger offspring sizes and live-bearing are a result of elevated predation in the tropics; given the body cavity constraint on offspring size of live-bearers we further speculate that large offspring size would require the evolution of larger adult body sizes in tropical rays that would allow greater maternal investment. This is consistent with the adult body size differences being the opposite of what might be expected under a TSR hypothesis, i.e. larger in tropical rays and smaller in cool-water skates, and may explain this exception to the TSR where tropical rays attain larger sizes at higher temperatures (Atkinson, 1995). Instead, our findings are more consistent with a mortality theory of life histories, and specifically the mortality arising from predation risk and offspring size (Auer *et al*., 2018; Glazier, 2023).

We found that offspring size, alongside temperature or depth, explained variation in *r*_max_ for rays and skates rather than adult body size or the adult-to-offspring size ratio. Offspring size reflects parental (mainly maternal) energy investment that is part of a key trade-off between reproduction, growth, and survival. Offspring size has direct fitness consequences through the allocation of resources provisioned to offspring for growth, survival, and reproductive success, and therefore has ecological and evolutionary implications (Pettersen, Schuster and Metcalfe, 2022). Given the strong phylogenetic signal in the *r*_max_ residuals, it is likely maximum population growth rate is shaped by biological traits that are evolutionary conserved, which would allow for predictive modelling of *r*_max_ based on phylogenetic relationships (Pardo and Dulvy, 2022). Additional variation in *r*_max_ that was not explained by our models may be explained by further environmental and physiological variables such as dissolved oxygen or metabolic rate (Pardo and Dulvy, 2022; Gravel *et al*., 2024). Indeed, further understanding of the relationship between offspring size and metabolic rate is needed (Pettersen, Schuster and Metcalfe, 2022). Empirical estimates of juvenile mortality for sharks and rays are still required to better understand juvenile survival across species with different life history strategies (incorporating how growth and mortality varies with body size). Whilst the mortality estimator used in the modified Euler-Lotka model to calculate *r*_max_ accounts for juvenile survival to maturity, there is a need for the development of size-dependent mortality rates to investigate differences across reproductive and offspring size strategies. Our results suggest that these differences may be key to understanding biogeographic patterns in extinction risk. We posit that greater predation risk in the tropics has driven the evolution of larger offspring size to increase offspring survival in tropical rays, potentially through live-bearing reproductive mode, increased matrotrophy, and larger adult body sizes. Consequently, shallow-water tropical rays have lower population growth rates and are more intrinsically sensitive to overfishing than may be expected from metabolic ecology.

## Acknowledgements

EB was funded by a Natural Environment Research Council (NERC) PhD studentship through the IAPETUS2 Doctoral Training Partnership (award ref. 2284959) and a UK Research and Innovation (UKRI)-Mitacs UK-Canada Globalink Research Placement (award ref. NE/T014555/1). JSB was supported by the National Science Foundation (NSF grant number 2109411). NKD was supported by the Natural Sciences and Engineering Research Council of Canada Discovery and Accelerator Awards (grant numbers 5013566 and 462291) and the Canada Research Chairs program (grant number 1228186).

## Data Availability Statement

The datasets supporting this article are available at FigShare (DOI: https://doi.org/10.6084/m9.figshare.20182109). The R code to reproduce these analyses will be available at Github (https://github.com/EBarrowclift/batoid-rmax-offspring-scaling).

## Conflict of Interest Statement

I confirm that all authors agree with the contents of the manuscript and its submission to Fish and Fisheries. All data and code sources have been made publicly available upon submission of the manuscript. Funding sources are fully acknowledged in the manuscript and the authors declare no competing interests.

## CRediT author contributions

Ellen Barrowclift: Conceptualisation, Methodology, Formal analysis, Investigation, Validation, Data curation, Writing-Original draft preparation, Writing-Review & Editing, Visualisation, Funding acquisition; Jennifer S. Bigman: Formal analysis, Writing-Reviewing and Editing; Eric D. Digel: Formal analysis, Writing-Reviewing and Editing; Per Berggren: Writing-Reviewing and Editing, Supervision, Funding acquisition; Nicholas K. Dulvy: Conceptualisation, Methodology, Formal analysis, Investigation, Validation, Data curation, Writing-Original draft preparation, Writing-Review & Editing, Visualisation, Funding acquisition, Supervision.

## Notes

https://doi.org/10.6084/m9.figshare.20182109

https://github.com/EBarrowclift/batoid-rmax-offspring-scaling

### Competing Interest Statement

The authors have declared no competing interest.

### Summary of Updates

Model clarification/simplification with updated supporting text in the introduction/methods/results/discussion.

## References

Arnold, T.W. (2010) ‘Uninformative parameters and model selection using Akaike’s Information Criterion’, The Journal of Wildlife Management, 74(6), pp. 1175–1178. Available at: 10.1111/j.1937-2817.2010.tb01236.x.

Ashton, G. V. et al. (2022) ‘Predator control of marine communities increases with temperature across 115 degrees of latitude’, Science, 376(6598), pp. 1215–1219. Available at: 10.1126/science.abc4916.

Atkinson, D. (1994) ‘Temperature and organism size-A biological law for ectotherms?’, Advances in Ecological Research, 25(C), pp. 1–58. Available at: 10.1016/S0065-2504(08)60212-3.

Atkinson, D. (1995) ‘Effects of temperature on the size of aquatic ectotherms: Exceptions to the general rule’, Journal of Thermal Biology, 20(1–2), pp. 61–74. Available at: 10.1016/0306-4565(94)00028-H.

Atkinson, D., Morley, S.A. and Hughes, R.N. (2006) ‘From cells to colonies: At what levels of body organization does the “temperature-size rule” apply?’, Evolution & development, 8(2), pp. 202–214. Available at: 10.1111/J.1525-142X.2006.00090.X.

Atkinson, D. and Sibly, R.M. (1997) ‘Why are organisms usually bigger in colder environments? Making sense of a life history puzzle’, Trends in ecology & evolution, 12(6), pp. 235–239. Available at: 10.1016/S0169-5347(97)01058-6.

Auer, S.K. et al. (2018) ‘Metabolic rate evolves rapidly and in parallel with the pace of life history’, Nature Communications, 9(14), p. 6. Available at: 10.1038/s41467-017-02514-z.

Barnes, C. et al. (2010) ‘Global patterns in predator-prey size relationships reveal size dependency of trophic transfer efficiency’, Ecology, 91(1), pp. 222–232. Available at: 10.1890/08-2061.1.

Barrowclift, E. et al. (2023) ‘Tropical rays are intrinsically more sensitive to overfishing than the temperate skates’, Biological Conservation, 281, p. 110003. Available at: 10.1016/j.biocon.2023.110003.

Barrowclift, E. and Dulvy, N.K. (2023) Life history and maximum weight database for rays (Class Chondrichthyes, Superorder Batoidea) up to June 2022, Figshare. Available at: 10.6084/m9.figshare.20182109.v3.

Bergmann, C. (1847) ‘Ueber die Verhältnisse der Wärmeökonomie der Thiere zu ihrer Grösse’, Göttinger Studien, 1, pp. 595–708.

Bernhardt, J.R., Sunday, J.M. and O’Connor, M.I. (2018) ‘Metabolic theory and the temperature-size rule explain the temperature dependence of population carrying capacity’, The American Naturalist, 192(6), pp. 687–697. Available at: 10.1086/700114.

Beukhof, E. et al. (2019) ‘Marine fish traits follow fast-slow continuum across oceans’, Scientific reports, 9(1), p. 17878. Available at: 10.1038/S41598-019-53998-2.

Blackburn, D.G. and Hughes, D.F. (2024) ‘Phylogenetic analysis of viviparity, matrotrophy, and other reproductive patterns in chondrichthyan fishes’, Biological Reviews, 99(4), pp. 1314–1356. Available at: 10.1111/BRV.13070.

Boettiger, C., Lang, D.T. and Wainwright, P.C. (2012) ‘rfishbase: Exploring, manipulating and visualizing FishBase data from R’, Journal of Fish Biology, 81(6), pp. 2030–2039. Available at: 10.1111/J.1095-8649.2012.03464.X.

Brown, J.H. et al. (2004) ‘Toward a metabolic theory of ecology’, Ecology, 85(7), pp. 1771– 1789. Available at: 10.1890/03-9000.

Burger, J.R., Hou, C. and Brown, J.H. (2019) ‘Toward a metabolic theory of life history’, Proceedings of the National Academy of Sciences of the United States of America, 116(52), pp. 26653–26661. Available at: 10.1073/PNAS.1907702116/SUPPL_FILE/PNAS.1907702116.SAPP.PDF.

Burnham, K.P. and Anderson, D.R. (2002) Model Selection and Multimodel Inference: A Practical Information-Theoretic Approach. 2nd edn, Springer. 2nd edn. Springer New York. Available at: 10.1007/b97636.

Conrath, C. and Musick, J. (2012) ‘Reproductive Biology of Elasmobranchs’, in J. Jeffrey C. Carrier, J.A. Musick, and M.R. Heithaus (eds) Biology of Sharks and Their Relatives. 2nd edn. Boca Raton, London, New York, Washington, D.C.: CRC Press, pp. 291–312. Available at: https://books.google.com/books/about/Biology_of_Sharks_and_Their_Relatives.html?id=7qXMBQAAQBAJ (Accessed: 27 January 2022).

Cortés, E. (2000) ‘Life history patterns and correlations in sharks’, Reviews in Fisheries Science, 8(4), pp. 299–344. Available at: 10.1080/10408340308951115.

Cortés, E. (2016) ‘Perspectives on the intrinsic rate of population growth’, Methods in Ecology and Evolution. Edited by J. Travis, 7(10), pp. 1136–1145. Available at: 10.1111/2041-210x.12592.

Denéchère, R., van Denderen, P.D. and Andersen, K.H. (2022) ‘Deriving population scaling rules from individual-level metabolism and life history traits’, The American Naturalist, 199(4), pp. 564–575. Available at: 10.1086/718642.

Duarte, C.M. and Alcaraz, M. (1989) ‘To produce many small or few large eggs: A size-independent reproductive tactic of fish’, Oecologia 1989 80:3, 80(3), pp. 401–404. Available at: 10.1007/bf00379043.

Dulvy, N.K. et al. (2014) ‘Extinction risk and conservation of the world’s sharks and rays’, eLife, 3, p. 590. Available at: 10.7554/eLife.00590.

Dulvy, N.K. et al. (2021) ‘Overfishing drives over one-third of all sharks and rays toward a global extinction crisis’, Current Biology, 31(21), pp. 4773-4787.e8. Available at: 10.1016/j.cub.2021.08.062.

Dulvy, N.K. and Reynolds, J. (1997) ‘Evolutionary transitions among egg-laying, live-bearing and maternal inputs in sharks and rays’, Proceedings of the Royal Society of London, B, 264, pp. 1309–1315.

Ebert, D.A. and Compagno, L.J.V. (2007) ‘Biodiversity and systematics of skates (Chondrichthyes: Rajiformes: Rajoidei)’, Environmental Biology of Fishes, 80(2–3), pp. 111– 124. Available at: 10.1007/S10641-007-9247-0/FIGURES/5.

Frisk, M. (2010) ‘Life History Strategies of Batoids’, in C. Carrier, J.A. Musick, and M.R. Heithaus (eds) Sharks and their Relatives II: Biodiversity, Adaptive Physiology, and Conservation. Boca Raton: CRC Press, pp. 283–316. Available at: 10.1201/9781420080483-c6.

Froese, R. and Pauly, D. (2022) FishBase. World Wide Web electronic publication. https://www.fishbase.org, (08/2022).

Glazier, D.S. (2023) ‘The relevance of time in biological scaling’, Biology, 12, p. 1084. Available at: 10.3390/biology12081084.

Goodwin, N.B., Dulvy, N.K. and Reynolds, J.D. (2002) ‘Life-history correlates of the evolution of live bearing in fishes’, Philosophical Transactions of the Royal Society of Biological Sciences, 357, pp. 259–267. Available at: 10.1098/rstb.2001.0958.

Gravel, S. et al. (2024) ‘Metabolism, population growth, and the fast-slow life history continuum of marine fishes’, Fish and Fisheries, 25, pp. 349–361. Available at: 10.1111/faf.12811.

Horswill, C. et al. (2025) ‘Imputation of fisheries reference points for endangered data-poor fishes, with application to Rhino Rays’, Fish and Fisheries, 0, pp. 1–18. Available at: 10.1111/faf.70003.

Hutchings, J.A. et al. (2012) ‘Life-history correlates of extinction risk and recovery potential’, Ecological Applications, 22(4), pp. 1061–1067. Available at: 10.1890/11-1313.1.

Jennings, S. et al. (2008) ‘Global-scale predictions of community and ecosystem properties from simple ecological theory’, Proceedings of the Royal Society B: Biological Sciences, 275(1641), pp. 1375–1383. Available at: 10.1098/RSPB.2008.0192.

Juan-Jordá, M.J. et al. (2011) ‘Global population trajectories of tunas and their relatives’, Proceedings of the National Academy of Sciences of the United States of America, 108(51), pp. 20650–20655. Available at: 10.1073/PNAS.1107743108.

Juan-Jordá, M.J. et al. (2013) ‘Life in 3-D: Life history strategies in tunas, mackerels and bonitos’, Reviews in Fish Biology and Fisheries, 23(2), pp. 135–155. Available at: 10.1007/S11160-012-9284-4/FIGURES/4.

Juan-Jordá, M.J. et al. (2015) ‘Population declines of tuna and relatives depend on their speed of life’, Proceedings of the Royal Society B: Biological Sciences, 282(1811). Available at: 10.1098/rspb.2015.0322.

Last, P.R. et al. (2016) Rays of the world. CSIRO Publishing.

Luhring, T.M. and Delong, J.P. (2017) ‘Scaling from metabolism to population growth rate to understand how acclimation temperature alters thermal performance’, Integrative and Comparative Biology, 57(1), pp. 103–111. Available at: 10.1093/icb/icx041.

Marshall, D.J. (2021) ‘Temperature-mediated variation in selection on offspring size: A multi-cohort field study’, Functional Ecology, 35(10), pp. 2219–2228. Available at: 10.1111/1365-2435.13879.

McEachran, J. and Miyake, T. (1990) Zoogeography and Bathymetry of skates (Chondrichthyes, Rajoidei), Elasmobranchs as living resources: advances in the biology, ecology, systematics, and the status of the fisheries. Edited by H.L. Pratt, S.H. Gruber, and T. Taniuchi. NOAA Technical Report National Marine Fisheries Service 90. Available at: https://www.semanticscholar.org/paper/Elasmobranchs-as-living-resources%3A-advances-in-the-Pratt-Gruber/b36e50b41efa176f12b596aa9ff55da8bbe5daef (Accessed: 14 April 2022).

Mejía, D. et al. (2025) ‘A global synthesis of population demographic models in sharks and rays’, Fish and Fisheries, 26(4), pp. 587–602. Available at: 10.1111/FAF.12900.

Moro, S. et al. (2025) ‘Living on the extinction edge: Resilience to fishing and rebound potential of the Mediterranean elasmobranchs’, Fish and Fisheries [Preprint]. Available at: 10.1111/FAF.12911.

Mull, C. et al. (2022) ‘Sharkipedia: A curated open-access database of shark and ray life history traits and abundance time-series’, Scientific Data., 9(559). Available at: 10.1038/s41597-022-01655-1.

Mull, C.G. et al. (2024) ‘Maternal investment evolves with larger body size and higher diversification rate in sharks and rays’, Current Biology, 34(12), pp. 2773-2781.e3. Available at: 10.1016/J.CUB.2024.05.019/ASSET/9293F57B-8BBE-414A-9832-002FD8F4431A/MAIN.ASSETS/GR4.JPG.

Musick, J.A. and Ellis, J.K. (2005) ‘Reproductive Evolution of Chondrichthyans’, in W.C. Hamlett (ed.) Reproductive Biology and Phylogeny of Chondrichthyes. 1st edn. Boca Raton: CRC press, pp. 45–78.

Myers, R.A., Mertz, G. and Fowlow, P.S. (1997) ‘Maximum population growth rates and recovery times for Atlantic cod, Gadus morhua’, Fishery Bulletin, 95, pp. 762–772.

Neuheimer, A.B. et al. (2015) ‘Adult and offspring size in the ocean over 17 orders of magnitude follows two life history strategies’, Ecology, 96(12), pp. 3303–3311. Available at: 10.1890/14-2491.1.

Olsson, K.H., Gislason, H. and Andersen, K.H. (2016) ‘Differences in density-dependence drive dual offspring size strategies in fish’, Journal of Theoretical Biology, 407, pp. 118–127. Available at: 10.1016/J.JTBI.2016.07.027.

Orme, D. et al. (2018) ‘caper: Comparative analyses of phylogenetics and evolution in R.’ Available at: https://cran.r-project.org/package=caper.

Pardo, S.A. et al. (2016) ‘Maximum intrinsic rate of population increase in sharks, rays, and chimaeras: The importance of survival to maturity’, Canadian Journal of Fisheries and Aquatic Sciences, 73(8), pp. 1159–1163. Available at: 10.1139/cjfas-2016-0069.

Pardo, S.A. and Dulvy, N.K. (2022) ‘Body mass, temperature, and depth shape the maximum intrinsic rate of population increase in sharks and rays’, Ecology and Evolution, 12, p. e9441. Available at: 10.1002/ece3.9441.

Pettersen, A.K. et al. (2020) ‘Linking life-history theory and metabolic theory explains the offspring size-temperature relationship’, Ecology Letters, 22(3), pp. 518–526. Available at: 10.1111/ele.13213.

Pettersen, A.K., Schuster, L. and Metcalfe, N.B. (2022) ‘The evolution of offspring size: A metabolic scaling perspective’, Integrative and Comparative Biology, 62(5), pp. 1492–1502. Available at: 10.1093/ICB/ICAC076.

R Core Team (2021) R, R: A Language and Environment for Statistical Computing. Vienna, Austria: R Foundation for Statistical Computing. Available at: https://www.r-project.org/.

RStudio Team (2021) ‘RStudio’, RStudio: Integrated Development Environment for R. [Preprint]. Boston, MA: RStudio, PBC. Available at: http://www.rstudio.com/.

Savage, V.M. et al. (2004) ‘Effects of body size and temperature on population growth’, The American Naturalist, 163(3), pp. 429–441. Available at: 10.1086/381872.

Sherman, S.C. et al. (2020) ‘When sharks are away, rays will play: Effects of top predator removal in coral reef ecosystems’, Marine Ecology Progress Series, 641, pp. 145–157. Available at: 10.3354/meps13307.

Sibly, R.M. et al. (2018) ‘The shark-tuna dichotomy: Why tuna lay tiny eggs but sharks produce large offspring’, Royal Society Open Science, 5(8), p. 180453. Available at: 10.1098/RSOS.180453.

Simpfendorfer, C.A. et al. (2023) ‘Widespread diversity deficits of coral reef sharks and rays’, Science, 380(6650), pp. 1155–1160. Available at: 10.1126/science.ade4884.

Smith, C.C. and Fretwell, S.D. (1974) ‘The optimal balance between size and number of offspring’, American Naturalist, 108(962), pp. 499–506. Available at: 10.1086/282929.

Stein, R.W. et al. (2018) ‘Global priorities for conserving the evolutionary history of sharks, rays and chimaeras’, Nature Ecology & Evolution 2018 2:2, 2(2), pp. 288–298. Available at: 10.1038/s41559-017-0448-4.

Thorson, J.T. (2020) ‘Predicting recruitment density dependence and intrinsic growth rate for all fishes worldwide using a data-integrated life-history model’, Fish and Fisheries, 21(2), pp. 237–251. Available at: 10.1111/FAF.12427.

Verity, P.G. and Smetacek, V. (1996) ‘Organism life cycles, predation, and the structure of marine pelagic ecosystems’, Marine Ecology Progress Series, 130, pp. 277–293.

Wong, S., Bigman, J.S. and Dulvy, N.K. (2021) ‘The metabolic pace of life histories across fishes’, Proceedings of the Royal Society B: Biological Sciences, 288(1953). Available at: 10.1098/rspb.2021.0910.

Wourms, J.P. (1994) ‘The challenges of piscine viviparity’, Israel Journal of Ecology and Evolution, 40(3–4), pp. 551–568. Available at: 10.1080/00212210.1994.10688772.

Wourms, J.P. and Lombardi, J. (1992) ‘Reflections on the evolution of piscine viviparity’, American Zoologist, 32(2), pp. 276–293. Available at: 10.1093/icb/32.2.276.

