## Supplementary Tables 1 - 4 for "Offspring size resolves a population growth paradox in rays and skates"

**Table S1:** All 32 models tested with associated hypotheses for how maximum intrinsic rate of population increase (*r*_max_) varies with adult body mass *M*, offspring body mass *M_offspring_*, inverse temperature $1/k_{B}T$, depth, and a temperature-depth index (PC1 axis from Principle Components Analysis of collapsed temperature and depth data in Barrowclift et al. (2023)). Comparison of 32 ln(*r*_max_) models using corrected Akaike Information Criteria (AICc), number of parameters (n), negative log-likelihood (-LL), R^2,^, adjusted R^2^ (Adj. R^2^), difference in AICc from the top model (ΔAICc), and Akaike weights. Models are ordered by ascending AICc, with models with AICc < 2 shown in bold. Note, Order was categorical for rays (Orders Myliobatiformes, Rhinopristiformes, and Torpediniformes) and skates (Order Rajiformes).

| **Rank** | **Hypothesis:** ***r*_max_ varies with** | **Model: ln(*r*_max_) ~** | **n** | **-LL** | **AICc** | **R^2^** | **Adj. R^2^** | **ΔAICc** | **Weights** |
| --- | --- | --- | --- | --- | --- | --- | --- | --- | --- |
| 1 | ***r*_max_ varies with offspring mass and depth** | **ln(*M_offspring_*) + depth** | 3 | -62.8 | 132.0 | 0.22 | 0.20 | 0.0 | 0.340 |
| 2 | ***r*_max_ varies with offspring mass and temperature** | **ln(*M_offspring_*) +** $\boldsymbol{1/}\boldsymbol{k}_{\boldsymbol{B}}\boldsymbol{T}$ | 3 | -63.0 | 132.3 | 0.21 | 0.19 | 0.3 | 0.293 |
| 3 | adult and offspring mass and temperature | ln(*M*) + ln(*M_offspring_*) + $1/k_{B}T$ | 4 | -63.0 | 134.4 | 0.21 | 0.18 | 2.4 | 0.103 |
| 4 | offspring mass only | ln(*M_offspring_*) | 2 | -65.4 | 135.0 | 0.17 | 0.16 | 3.0 | 0.076 |
| 5 | adult and offspring mass | ln(*M*) + ln(*M_offspring_*) | 3 | -65.3 | 136.9 | 0.17 | 0.15 | 4.9 | 0.029 |
| 6 | adult mass and temperature-depth index | ln(*M*) + temperature-depth index | 3 | -65.4 | 137.2 | 0.16 | 0.14 | 5.2 | 0.025 |
| 7 | adult and offspring mass, and the effect of mass scaling coefficient varies with offspring size | ln(*M*) * ln(*M_offspring_*) | 4 | -64.9 | 138.2 | 0.17 | 0.14 | 6.2 | 0.015 |
| 8 | adult mass, temperature-depth index, and Order | ln(*M*) + temperature-depth index + Order | 4 | -65.0 | 138.4 | 0.17 | 0.14 | 6.4 | 0.014 |
| 9 | adult mass and depth | ln(*M*) + depth | 3 | -66.1 | 138.6 | 0.14 | 0.12 | 6.6 | 0.013 |
| 10 | adult mass and temperature | ln(*M*) + $1/k_{B}T$ | 3 | -66.1 | 138.5 | 0.14 | 0.12 | 6.5 | 0.013 |
| 11 | adult mass and the effect of temperature varies with depth | ln(*M*) + $1/k_{B}T$ * depth | 5 | -64.1 | 138.9 | 0.18 | 0.14 | 6.9 | 0.011 |
| 12 | adult mass and temperature-depth index, and the effect of mass scaling coefficient varies with the temperature-depth index | ln(*M*) * temperature-depth index | 4 | -65.3 | 139.1 | 0.16 | 0.13 | 7.1 | 0.010 |
| 13 | adult mass, temperature, and depth | ln(*M*) + $1/k_{B}T$ + depth | 4 | -65.4 | 139.3 | 0.16 | 0.13 | 7.3 | 0.009 |
| 14 | adult mass and temperature, and the effect of mass scaling coefficient varies with temperature | ln(*M*) * $1/k_{B}T$ | 4 | -65.7 | 139.8 | 0.16 | 0.13 | 7.8 | 0.007 |
| 15 | adult mass, temperature, and Order | ln(*M*) + $1/k_{B}T$ + Order | 4 | -65.6 | 139.7 | 0.15 | 0.12 | 7.7 | 0.007 |
| 16 | adult mass only | ln(*M*) | 2 | -68.0 | 140.2 | 0.10 | 0.09 | 8.2 | 0.006 |
| 17 | adult mass, depth, and Order | ln(*M*) + depth + Order | 4 | -65.9 | 140.2 | 0.15 | 0.12 | 8.2 | 0.006 |
| 18 | adult mass, temperature-depth index, and Order, and the effect of mass scaling coefficient varies with the temperature-depth index | ln(*M*) * temperature-depth index + Order | 5 | -64.8 | 140.4 | 0.17 | 0.13 | 8.4 | 0.005 |
| 19 | adult mass and depth, and the effect of mass scaling coefficient varies with depth | ln(*M*) * depth | 4 | -66.1 | 140.8 | 0.14 | 0.11 | 8.8 | 0.004 |
| 20 | adult mass, temperature, and Order, and the effect of mass scaling coefficient varies with temperature | ln(*M*) * $1/k_{B}T$ + Order | 5 | -65.1 | 141.0 | 0.17 | 0.13 | 9.0 | 0.004 |
| 21 | adult mass and Order | ln(*M*) + Order | 3 | -68.0 | 142.2 | 0.11 | 0.08 | 10.2 | 0.002 |
| 22 | adult mass, depth, and Order, and the effect of mass scaling coefficient varies with depth | ln(*M*) * depth + Order | 5 | -65.9 | 142.5 | 0.15 | 0.11 | 10.5 | 0.002 |
| 23 | temperature-depth index only | temperature-depth index | 2 | -69.1 | 142.3 | 0.08 | 0.07 | 10.3 | 0.002 |
| 24 | depth only | depth | 2 | -70.3 | 144.8 | 0.05 | 0.04 | 12.8 | 0.001 |
| 25 | temperature only | $1/k_{B}T$ | 2 | -69.5 | 143.1 | 0.07 | 0.06 | 11.1 | 0.001 |
| 26 | temperature and Order | $1/k_{B}T$ + Order | 3 | -68.8 | 144.0 | 0.09 | 0.06 | 12.0 | 0.001 |
| 27 | temperature-depth index and Order | temperature-depth index + Order | 3 | -68.5 | 143.3 | 0.09 | 0.07 | 11.3 | 0.001 |
| 28 | adult:offspring size ratio and temperature | ln(*M*/*M_offspring_*) + $1/k_{B}T$ | 3 | -69.2 | 144.8 | 0.08 | 0.06 | 12.8 | 0.001 |
| 29 | average *r*_max_ (i.e. intercept-only model) | 1 | 1 | -72.7 | 147.5 | 0.00 | 0.00 | 15.5 | 0.000 |
| 30 | Order | 1 + Order | 2 | -72.6 | 149.4 | 0.00 | -0.01 | 17.4 | 0.000 |
| 31 | depth and Order | depth + Order | 3 | -70.1 | 146.4 | 0.06 | 0.04 | 14.4 | 0.000 |
| 32 | adult:offspring size ratio only | ln(*M*/*M_offspring_*) | 2 | -72.7 | 149.5 | 0.00 | -0.01 | 17.5 | 0.000 |

**Table S2: Sensitivity test for the choice of phylogenetic tree.** Comparison of 11 ln(*r*_max_) models fitted with 10 different phylogenetic trees obtained from Stein et al. (2018) (available on Vertlife.org) using corrected Akaike Information Criteria (AICc). The model with lowest AICc for each iteration is shown in bold.

| **ln(*r*_max_) ~** | **1** | **2** | **3** | **4** | **5** | **6** | **7** | **8** | **9** | **10** |
| --- | --- | --- | --- | --- | --- | --- | --- | --- | --- | --- |
| **ln(*M_offspring_*) + depth** | 0.8 | **0** | **0** | 0.2 | **0** | 0.3 | 0.3 | **0** | 0.2 | 0.1 |
| **ln(*M_offspring_*) +** $\boldsymbol{1/}\boldsymbol{k}_{\boldsymbol{B}}\boldsymbol{T}$ | **0** | 1.5 | **0** | **0** | 0.5 | **0** | **0** | 1.1 | **0** | **0** |
| ln(*M_offspring_*) | 2.7 | 3.4 | 2.8 | 3.1 | 3.2 | 2.4 | 2.1 | 4.1 | 2.3 | 3.3 |
| ln(*M*) + $1/k_{B}T$ | 6.9 | 10.1 | 6.6 | 7.1 | 10.5 | 10.5 | 7.4 | 7.3 | 10.7 | 8 |
| ln(*M*) + depth | 7 | 9 | 6.8 | 7.4 | 9.8 | 11 | 7.7 | 6.1 | 10.8 | 8.9 |
| ln(*M*) * $1/k_{B}T$ | 8.7 | 11.9 | 7.7 | 9.3 | 12.6 | 12.7 | 9.2 | 9.2 | 12.8 | 9.9 |
| ln(*M*) | 8.5 | 10.6 | 8.5 | 9.5 | 12.2 | 12.1 | 8.4 | 9.2 | 12.4 | 10.4 |
| ln(*M*) * depth | 9.2 | 11 | 9 | 14.7 | 11.5 | 12.6 | 9.9 | 8.3 | 13 | 10.9 |
| $1/k_{B}T$ | 11.1 | 14.6 | 9.1 | 12 | 14.4 | 14 | 11 | 11.9 | 14.6 | 11.9 |
| depth | 13 | 15 | 10.8 | 14.2 | 14.9 | 15.6 | 12.8 | 12 | 15.5 | 14.3 |
| 1 | 15.1 | 18 | 13.5 | 16.8 | 17.9 | 17.4 | 14.4 | 16.3 | 18.1 | 16.6 |

**Table S3:** **Sensitivity test for the exclusion of manta rays.** Comparison of 11 ln(*r*_max_) models fitted without data for two manta ray species (*Mobula alfredi* and *M. birostris*) with largest offspring sizes using corrected Akaike Information Criteria (AICc), number of parameters (n), negative log-likelihood (-LL), R^2,^, adjusted R^2^ (Adj. R^2^), difference in AICc from the top model (ΔAICc), and Akaike weights. Models are ordered by ascending AICc and models with AICc < 2 shown in bold.

| **ln(*r*_max_) ~** | **n** | **-LL** | **AICc** | **R^2^** | **Adj. R^2^** | **ΔAICc** | **Weights** |
| --- | --- | --- | --- | --- | --- | --- | --- |
| **ln(*M_offspring_*) + depth** | **3** | **-59** | **124.3** | **0.15** | **0.13** | **0** | **0.382** |
| **ln(*M_offspring_*) +** $\boldsymbol{1/}\boldsymbol{k}_{\boldsymbol{B}}\boldsymbol{T}$ | **3** | **-59.3** | **124.8** | **0.14** | **0.12** | **0.5** | **0.297** |
| ln(*M_offspring_*) | 2 | -61.2 | 126.5 | 0.1 | 0.09 | 2.2 | 0.127 |
| ln(*M*) + depth | 3 | -61 | 128.2 | 0.1 | 0.08 | 3.9 | 0.054 |
| ln(*M*) + $1/k_{B}T$ | 3 | -61.1 | 128.5 | 0.1 | 0.08 | 4.2 | 0.047 |
| ln(*M*) | 2 | -62.7 | 129.6 | 0.07 | 0.05 | 5.3 | 0.027 |
| ln(*M*) * depth | 4 | -61 | 130.4 | 0.11 | 0.07 | 6.1 | 0.018 |
| ln(*M*) * $1/k_{B}T$ | 4 | -60.9 | 130.4 | 0.11 | 0.08 | 6.1 | 0.018 |
| $1/k_{B}T$ | 2 | -63.4 | 131 | 0.05 | 0.04 | 6.7 | 0.013 |
| depth | 2 | -63.6 | 131.4 | 0.04 | 0.03 | 7.1 | 0.011 |
| 1 | 1 | -65.5 | 133 | 0 | 0 | 8.7 | 0.005 |

**Table S4:** Coefficient estimates (95% confidence intervals estimated from standard errors shown in brackets) for all 11 models of ln(*r*_max_). Models are ordered by ascending AICc, with models with ∆AICc < 2 shown in bold. Pagel’s λ indicates the strength of the phylogenetic signal.

| **ln(*r*_max_) ~** | **intercept** | **ln(*M*)** | **depth** | $\boldsymbol{1/}\boldsymbol{k}_{\boldsymbol{B}}\boldsymbol{T}$ | **ln(*M*):**  **depth** | **ln(*M*):**  $\boldsymbol{1/}\boldsymbol{k}_{\boldsymbol{B}}\boldsymbol{T}$ | **ln(*M_offspring_*)** | **Pagel's λ** |
| --- | --- | --- | --- | --- | --- | --- | --- | --- |
| **ln(*M_offspring_*) + depth** | **-0.56**  **(-1.09 , -0.03)** | **-** | **-0.32**  **(-0.59 , -0.05)** | **-** | **-** | **-** | **-0.17**  **(-0.25 , -0.09)** | **0.79**  **(0.48 , 0.92)** |
| **ln(*M_offspring_*) +** $\boldsymbol{1/}\boldsymbol{k}_{\boldsymbol{B}}\boldsymbol{T}$ | **-0.62**  **(-1.16 , -0.08)** | **-** | **-** | **-0.44**  **(-0.82 , -0.05)** | **-** | **-** | **-0.16**  **(-0.24 , -0.08)** | **0.80**  **(0.49 , 0.93)** |
| ln(*M_offspring_*) | **-0.54**  **(-1.09 , 0.01)** | **-** | **-** | **-** | **-** | **-** | **-0.17**  **(-0.26 , -0.09)** | 0.80  (0.52, 0.93) |
| ln(*M*) + $1/k_{B}T$ | -0.23  (-1.13 , 0.68) | -0.11  (-0.19 , -0.03) | - | -0.41  (-0.83 , 0) | - | - | - | 0.89  (0.68, 0.96) |
| ln(*M*) + depth | -0.11  (-0.99 , 0.78) | -0.12   (-0.19 , -0.04) | -0.27  (-0.55 , 0) | - | - | - | - | 0.88  (0.67, 0.96) |
| ln(*M*) * $1/k_{B}T$ | -0.2   (-1.13 , 0.73) | -0.11  (-0.19 , -0.03) | - | -1.16  (-2.61 , 0.29) | - | 0.07  (-0.06 , 0.21) | - | 0.92  (0.72, 0.98) |
| ln(*M*) | -0.01  (-0.91 , 0.89) | -0.12   (-0.2 , -0.05) | - | - | - | - | - | 0.88  (0.69, 0.96) |
| ln(*M*) * depth | -0.1   (-1 , 0.79) | -0.12  (-0.2 , -0.04) | -0.3   (-1.75 , 1.15) | - | 0  (-0.15 , 0.16) | - | - | 0.88  (0.63, 0.96) |
| $1/k_{B}T$ | -1.22  (-1.76 , -0.68) | - | - | -0.55   (-0.96 , -0.13) | - | - | - | 0.89  (0.71, 0.97) |
| depth | -1.18  (-1.71 , -0.65) | - | -0.32   (-0.6 , -0.03) | - | - | - | - | 0.88  (0.68, 0.96) |
| 1 | -1.17  (-1.71 , -0.62) | - | - | - | - | - | - | 0.88  (0.69, 0.96) |
